# MIDOE: Maximally-informed Design of Experiment to infer experimentally inaccessible transcription factors dynamics

**DOI:** 10.1101/2025.08.04.668461

**Authors:** Razeen Shaikh, Gregory T. Reeves

## Abstract

The TGF-β/Smad signaling pathway regulates growth, development, and homeostasis of tissues across the animal kingdom. The pathway is activated when transforming growth factor -β (TGF-β) binds to its cognate transmembrane receptors to activate Smad2 by phosphorylation. The activated phosphor-Smad2 (PSmad2) undergoes a series of biochemical interactions with transcriptional activator Smad4 to form several oligomers, including the transcription factor (PSmad2)_2_/Smad4, which regulates target gene expression. Quantitative live cell imaging and mathematical modeling have been used to estimate the dynamics of (PSmad2)_2_/Smad4. However, due to the emergent nature of Smad2-Smad4 interactions, deconvolving the dynamics of the (PSmad2)_2_/Smad4 is challenging. We show that the Smad model is sloppy, has large parameter uncertainties (Ο∼10^15^), and the eigenvalues of the Fisher Information Matrix span over several decades. As such, well-fit parameter sets generate highly under-constrained predictions. To overcome this, we employ Profile Log-Likelihood to guide maximally informed design of experiments (MIDOE) and infer the dynamics of (PSmad2)_2_/Smad4. We generate these experiments computationally and validate that MIDOE can optimally constrain model predictions. We demonstrate that such careful analysis would not only improve the predictive power of models in systems biology but also reduce the time and expense of performing non-optimal experiments.

## 2 INTRODUCTION

Mathematical models are widely applied in systems biology to summarize signal transduction pathways, including the TGF-β/Smad pathway (**Figure 1A**), design experiments, and predict the effect of perturbation on system behavior. These are typically ODE-based deterministic models formulated to describe the rate of change in species as a function of time and variable model parameters (model structure), including rate constants and initial concentrations. The variable parameters are estimated by solving the inverse problem of fitting the model to the data and evaluating goodness-of-fit metrics (χ^2^) (**Figure 1B**).

**Figure 1:**
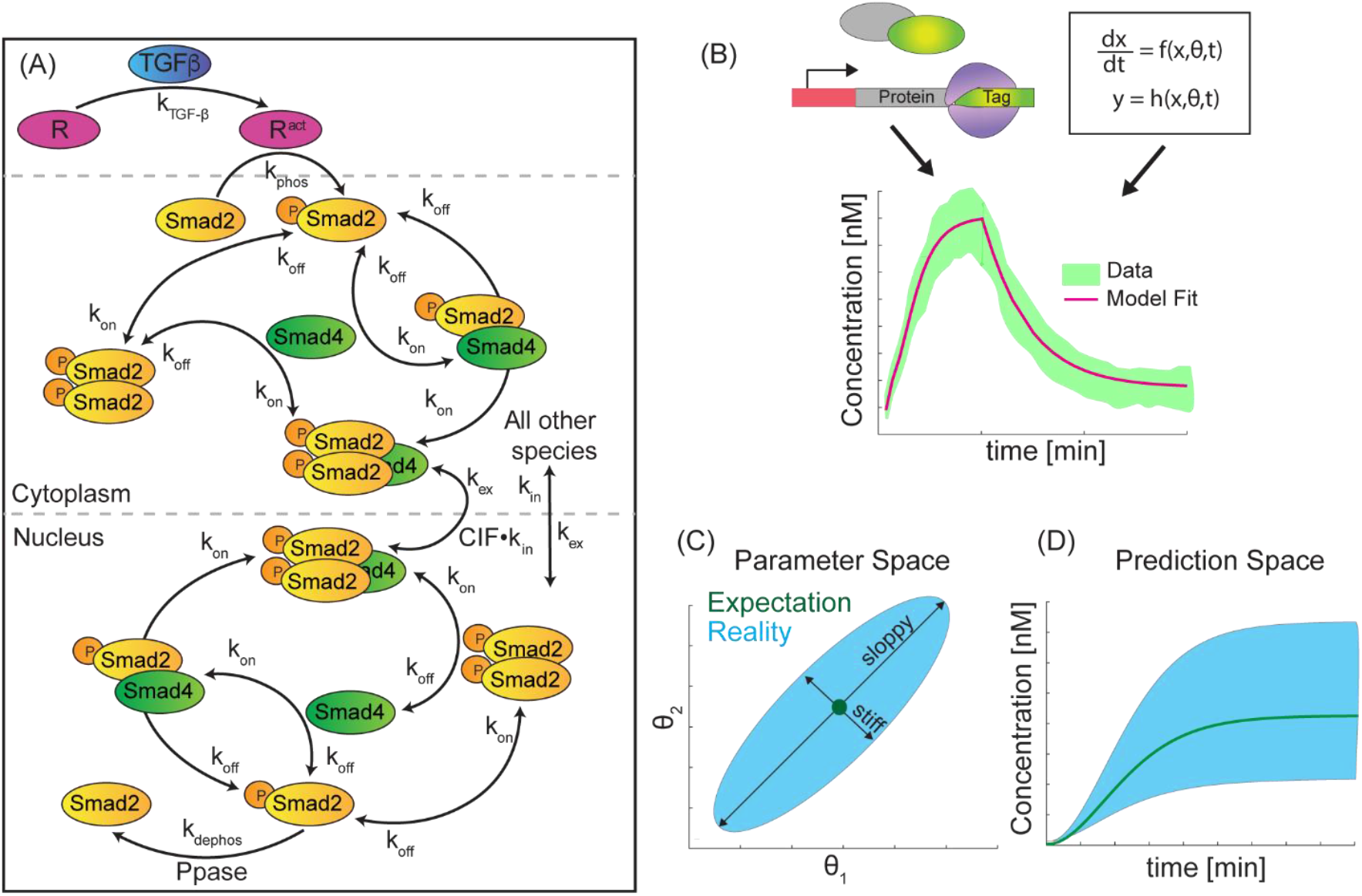
Systems Biology models are sloppy. **(A)** The TGF-β/Smad signal transduction network. **(B-D)** An illustration of a two-parameter fit to experimental data, which results in “well-fit” model outcomes **(B)**; however, the inferred parameters lie on a manifold characterized by a long (sloppy) axis and a narrow (stiff) axis **(C)**, resulting in under-constrained model predictions **(D)**

However, the inferred parameters have large uncertainties associated with them such that they are not point values; rather, they are parameter clouds (**Figure 1C**)^1^. The distributions of these “well-fit” parameter sets are within a confidence threshold of χ^2^, and can span over several orders of magnitude in parameter space^2,3^. These parameter clouds represent the feasible space of solutions, and their shape and size depend on the model topology and the experimental data. Each parameter set in the cloud fits the data comparably well, and one cannot distinguish between one parameter set and another.

To make matters worse, model predictions stemming from different ends of the parameter cloud are highly under-constrained, diminishing the utility of the model (**Figure 1D**). As such, it is important to ascertain whether the model parameters can be accurately estimated from the information present in the data. Parameters that can be estimated under the given model structure and data are deemed *identifiable*, whereas those that cannot be accurately estimated are *unidentifiable*. This aids the design of experiments (DOE), which will be maximally informative in constraining the remaining parameters.

The uncertainties in predictions associated with these models arise from χ^2^ and are characterized by the eigenvalues (λ) of a Fisher Information Matrix (FIM)^2,4-6^ — a measure of the amount of information an observable random variable carries about an unknown parameter. Typically, the distribution of eigenvalues spans over many decades, such that large uncertainties in parameter estimates correspond to wide distribution in eigenvalues^7^. Geometrically, for a mathematical model with *N* parameters and *M* data points, the model manifold is a *N*-dimensional object in *M*-dimensional space, such that the width of each axis of the object (*w*_*i*=1,...,N_) is inversely proportional to the square root of the *λ*_*i*=1,…,N_. Models in systems biology have a large number of parameters and are typically calibrated with sparse data, which renders a long and thin hyperribbon-like structure to the model manifold^7,8^. Such models are deemed *sloppy*.

One approach to reducing the uncertainty in parameter estimates is to perform more experiments and gather more information about system behavior. However, further experiments can be expensive and time-consuming, and the information gathered through these experiments may not be sufficient to accurately estimate parameters and make reliable predictions^1^. To effectively calibrate models, we need to design a set of maximally-informative experiments to reliably constrain the predictions. To address these issues, we use Profile Log-Likelihood (PLL), which is a technique to systematically characterize the identifiability of parameters. PLL evaluates the range over which a given parameter can be estimated from the available data and leads to a, rationale design of maximally informative experiments^3,9,10^.

In this manuscript, we detail a general procedure to reduce parameter uncertainty, and thereby increase the predictiveness of systems biology models. As an example, we focus on the TGF-β/Smad signal transduction pathway. The TGF-β/Smad signal transduction pathway regulates growth, development, and homeostasis of tissues and is conserved at a molecular and functional level from worms to humans^11-14^. The pathway is activated when transforming growth factor-β (TGF-β), an extracellular ligand from the TGF-β super-family, binds to the cognate transmembrane receptors. The ligand-receptor complex transduces the signal intracellularly by phosphorylating Smad2. Phospho-Smad2 (PSmad2) takes part in a series of biochemical interactions with transcriptional activator Smad4 to form several homomeric and heteromeric species, including the heterotrimeric transcription factor (PSmad2)_2_/Smad4, which translocates to the nucleus to regulate target gene expression.

Due to the complex nature of the Smad network topology, mathematical models have been deployed to infer the intracellular dynamics of the pathway. Live cell imaging of an EGFP-tagged Smad2 cell line, together with inverse modeling, has been used to estimate the dynamics of (PSmad2)_2_/Smad4^14,15^. However, due to the emergent dynamics of Smad2 and Smad4 interactions, deconvolving the dynamics of (PSmad2)_2_/Smad4 is challenging. To overcome these challenges, we first show that the Smad model fit to RFP-Smad2 data is sloppy, which means there is a broad spectrum of “well-fit” parameter sets. As such, the model has large parameter uncertainties (Ο∼10^15^), and the eigenvalues of the FIM span over fifteen decades. Consequently, we show that the spectrum of well-fit parameter sets generates highly under-constrained predictions. To constrain the Smad model predictions, we employed Profile Log-Likelihood (PLL) analysis to determine the range over which each parameter cannot be learned from data — parametric and structural unidentifiabilities. Next, using the results from the PLL analysis, we simulated a series of feasible experiments to determine which experimental design would result in maximally informed parameter estimation and in highly constrained predictions. Finally, we used further literature data to show that maximally informed design of experiment (MIDOE) results in a model with reduced uncertainty, increased parameter identifiability, and highly constrained predictions. We suggest that these approaches could be used to inform the study of a wide variety of biological systems.

## 3 Results

### The Dynamics OF THE Smad Signaling Complex ARE Under-constrained

We adapted a model of the Smad pathway which was formulated to account for the various known interactions in the canonical Smad pathway^16^. In particular, the model formulation includes Smad2 and Smad4 hetero- and homo-dimers, which interact to form the Smad signaling complex (SC): (PSmad2)_2_/Smad4^11,17,18^. We calibrated this model with RFP-Smad2 dynamics in C2Cl2 cells^19^, hereafter referred to as “Smad_v0” model. The data include the nuclear localization dynamics of RFP-Smad2 upon pathway activation by TGF-β, followed by depletion after treatment with SB-431542, a potent and specific inhibitor of the TGF-β pathway. It should be noted that these fluorescence measurements lump together all Smad2-containing species in the nuclei, including phosphor-forms, homo- and heterodimers, as well as the hetero-tetrameric signaling complex (SC). We used an evolutionary algorithm, *ISRES+*^20^, to fit the Smad model to the RFP-Smad2 data by evaluating an objective function (χ^2^) formulated as the sum of the squared errors (SSE) between the model outcomes and experimental data, normalized by experimental error. We repeated this approach several times (N = 89) and observed that the range of the distribution of inferred parameter estimates has an order of magnitude (*O*) of seven (**Figure 2A**). The inferred parameter sets were positive-definite and hence optimal solutions to the coupled ODEs. While these parameter sets were all well-fit, in that they each faithfully reproduced the RFP-Smad2 traces within experimental error (**Figure 2B**), they resulted in distinctive SC predictions (**Figure 2C**). For instance, the predictions of SC dynamics for an arbitrarily chosen parameter set is indicated in magenta (**Figure 2A,C**). We summarized the variability in the dynamics of SC by quantifying its dynamic characteristics, including dead time, time constant, rise time, steady state concentration of SC and fractional overshoot (**Figure 2C**). The histograms for the dynamic characteristics of SC indicate that the Smad model predictions span across a wide range with a high coefficient-of-variation (CV = σ/μ) (**Figure 2D-H**).

**Figure 2:**
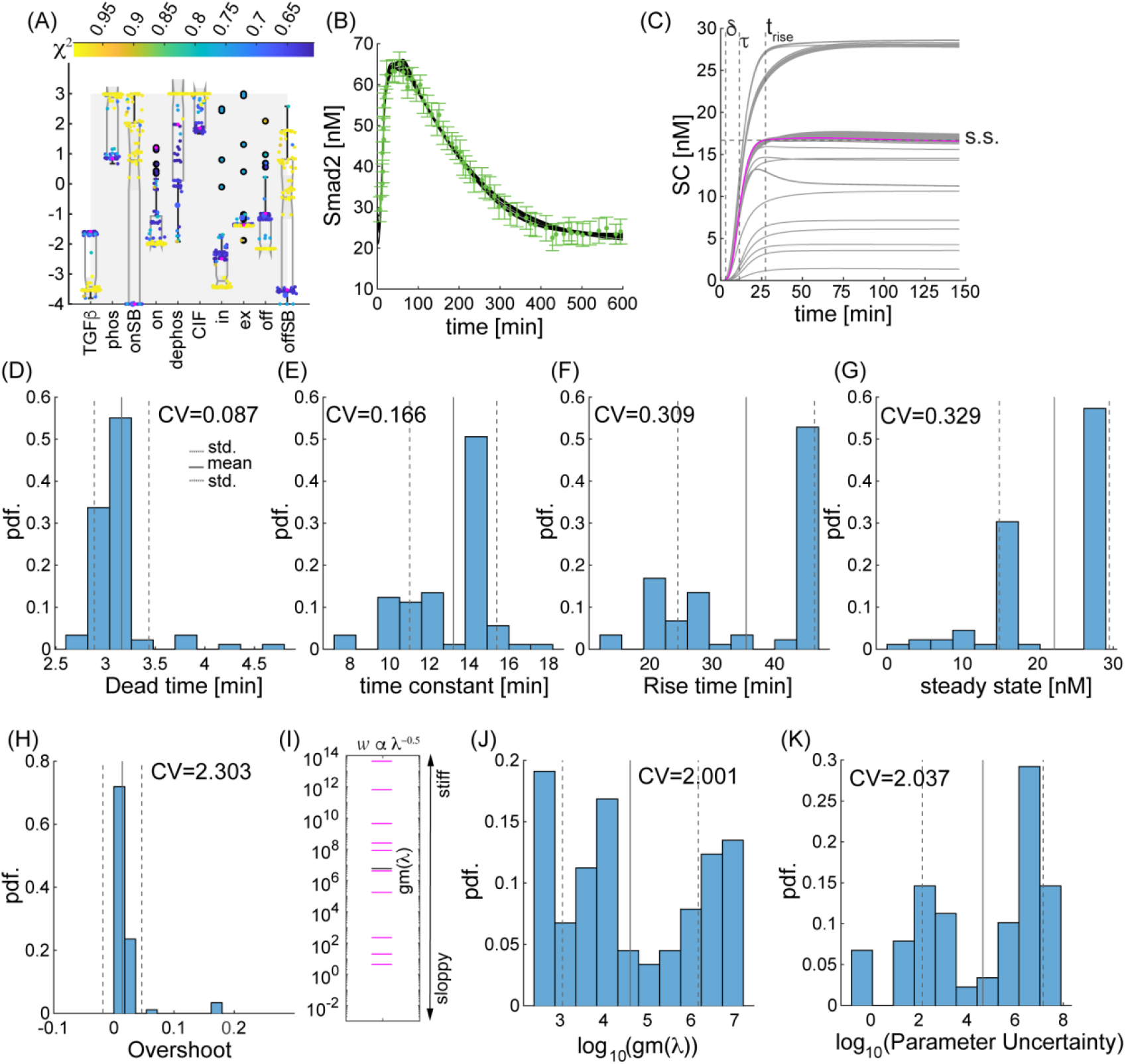
The predictions of the Smad signaling model are under-constrained. **(A-C)** The distribution of inferred parameter sets **(A)** for the Smad signaling model calibrated with Smad2 experimental data (green, ±SD) results in excellent fits (black) **(B)** but under-constrained SC dynamics **(C)**. In (A,C), a representative parameter set was arbitrarily chosen for concrete illustration purposes (magenta color). **(D-H)** The histograms of the dynamic characteristics of the SC, including dead time (δ) **(D)**, time constant (τ) **(E)**, rise time (t_rise_) **(F)**, steady state (s.s.) concentration of SC **(G)** and fractional overshoot **(H)**, span across a wide range. Several of these dynamic characteristics are illustrated in (C) for the representative parameter set (magenta). **(I)** The eigenvalues of the FIM of representative parameter set in (A, magenta) and (C, magenta). gm(λ): geometric mean of the eigenvalues. **(J-K)** Model calibration metrics including geometric mean of the eigenvalues of the FIM **(J)** and parameter uncertainty **(K)** range over several orders of magnitude, a characteristic feature of sloppy models.

We evaluated the extent of model calibration by computing the spectrum of the eigenvalues of the Fisher-Information Matrix (FIM), a measure of the amount of information an observable random variable carries about an unknown parameter^2,4-6^. For the parameter set visualized in magenta, the distribution of eigenvalues of the FIM span fourteen decades (**Figure 2I**). A separation of more than three orders of magnitude is a characteristic of a sloppy model^5^. The geometric mean of the eigenvalues of the FIM (*gm(*λ_FIM_)) (**Figure 2I**; black dash) for the Smad signaling model range over five orders of magnitude (**Figure 2J**). Another metric to evaluate the model calibration using FIM is Parameter Uncertainty (PU). Theoretically, near the best-fit solution, there is a set of “well-fit” solutions that, while not optimal, still result in model fits that align with the data within the limits of experimental noise. This region defines the confidence interval for the parameter estimate. The corresponding variation in parameters is defined as PU, which can be estimated from the FIM^2^ as:

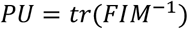

The range of the distribution of PU for all the optimal parameter sets of the Smad signaling model is over ten orders of magnitude (**Figure 2K**).

The signaling complex SC is a key response variable in the TGF-β signal transduction pathway and accurately predicting its dynamics is crucial^11,13^. To achieve this goal, we employed maximally informed design of experiments (MIDOE) methodology to constrain the variability in the predictions of the SC.

### The Smad SIGNALING MODEL HAS HIGHLY VARIABLE PREDICTIONS ALONG THE SLOPPY DIRECTIONS

The variability in model predictions could arise due to either the presence of multiple local minima (i.e., due to the model structure^1,10,21^) or due to a broad spectrum of well-fit parameter sets around a single minimum. Geometrically, the width (*w*) of the hyperribbon-shaped model manifold along a given eigenvector is inversely proportional to the square root of the corresponding eigenvalue (λ_FIM_), such that the lowest eigenvalues result in sloppiest widths^2,7,22^ (**Figure 2I**). We reconstructed a *pseudo-model manifold* around each inferred parameter set by sampling within a three-dimensional ellipsoid whose axes were approximated from the three sloppiest eigenvectors of the FIM, within confidence threshold. The (point-wise) confidence threshold (CT) is defined as the log-likelihood of the best fit parameter set plus the 99%-quantile of the χ^2^-distribution with degrees of freedom equal to number of parameters (see Methods). We then sampled at least a thousand parameter sets within the *pseudo-model manifold* around each parameter estimate and evaluated their goodness-of-fit metrics and model predictions. Several of the sampled parameter sets resulted in high goodness-of-fit (low χ^2^) but there were a few base parameters sets for which the sampled sets either included negative parameters or resulted in absurdly poor fits (high χ^2^). We reasoned that this was because the *pseudo-model manifold* is an approximation of the actual model manifold and, in some cases, does not align with the actual model manifold (**Figure 3D**). For instance, the *pseudo-model manifold* for the parameters visualized in magenta in the previous section (**Figure 2A**), was created by sampling at least a thousand parameter sets around the base parameter set, such that the sampling space was bound by an ellipsoid whose axes were estimated with eigendirections of the three sloppiest (smallest λ_FIM_) eigenvalues within confidence intervals. To aid visualization of this *pseudo-model manifold*, we used *t-SNE* to represent the parameter space in two dimensions (**Figure 3A**). Note that, while the parameter sets sampled around the reference parameter set result in high goodness-of-fits (**Figure 3B**), the predictions of the SC (although not as under-constrained as in **Figure 2C**) have a wide spectrum of dynamic characteristics, as measured by their CV (**Figure 3C**). We created the *pseudo-model manifold* around each inferred parameter set (**Figure 2A**) and computed the CV of the characteristics of the SC dynamics (**Figure 3E**). The ratio of the CV of the sampled sets to the CV for Smad_v0 ranges between 10-90% (**Figure 3F**). Based on this analysis, we inferred that, while the existence of multiple local minima contributes to the variability in the predictions of the Smad model, the majority of the variability in predictions arises from the sloppiness of model calibration.

**Figure 3:**
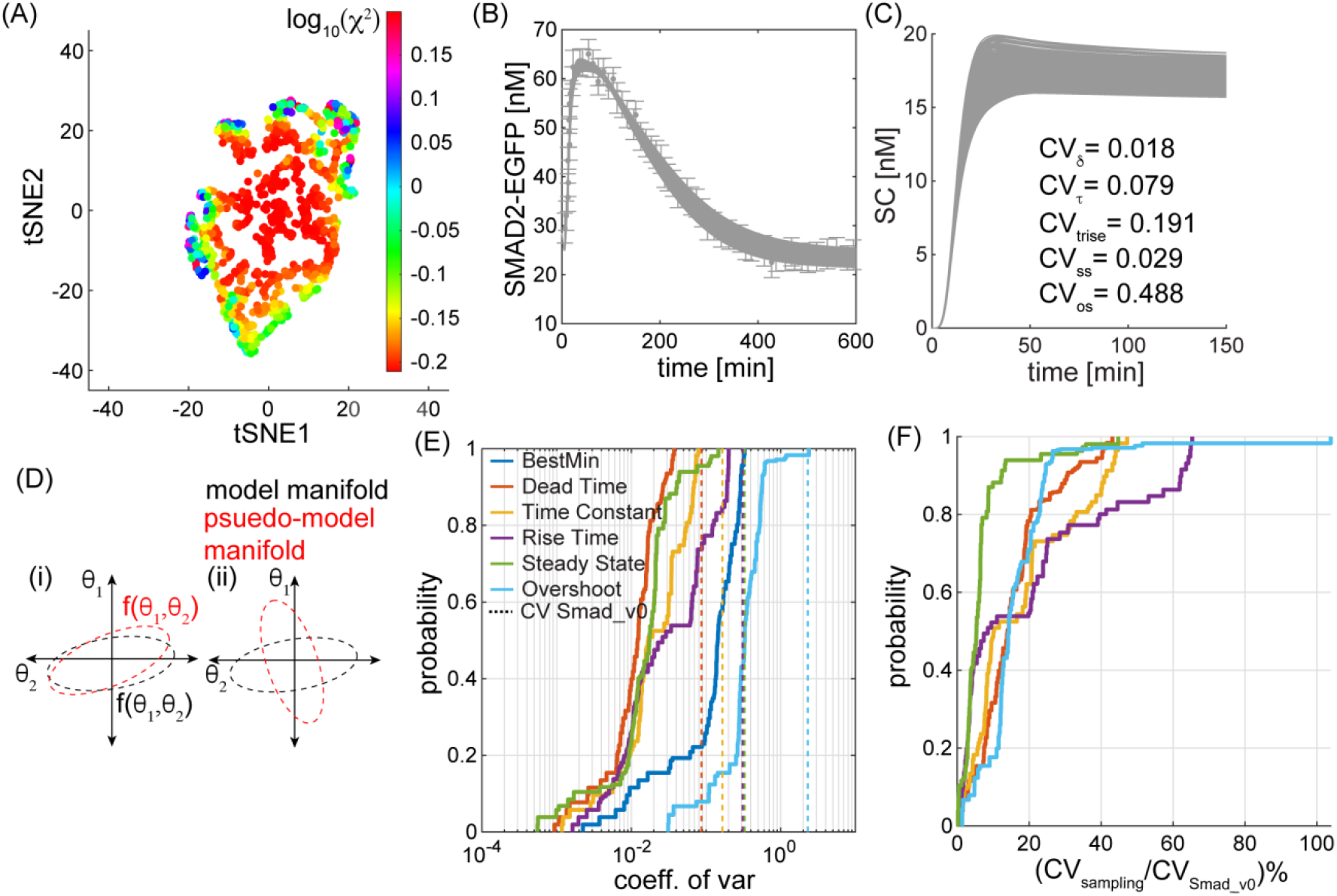
pseudo-Model manifolds along sloppy eigendirections. **(A)** t-SNE reconstruction of an ellipsoidal space bounded by confidence threshold of 1.6. **(B-C)** The Smad model fits **(B)** and predictions **(C)** for the sampled non-negative parameter sets within the ellipsoid for which the objective function is less than confidence threshold of 1.6. **(D)** An illustration of two cases **(i)** pseudo-model manifold aligns with the actual model manifold and **(ii)** the pseudo-model manifold is oriented such that it does not align with the actual manifold. **(E-F)** The probability distribution of the coefficient of variation **(E)** and ratio of CV of sampled sets over the CV of Smad_v0 **(F)** computed across all the feasible parameters set in each pseudo-Model manifold.

### Experimental DESIGN WITH Parameter IDENTIFIABILITY THROUGH LIKELIHOOD ESTIMATION

The Smad model is sloppy due to a broad spectrum of well-fit parameter sets around a single minimum, which resulted in under-constrained parameter estimates. One approach to reducing the sloppiness could be to measure the exact value of each parameter *in vivo*. However, this is expensive, likely infeasible, and has been shown to not result in accurate model predictions^1,23^.To address these issues, we adopt a parameter identifiability approach, Profile Log-Likelihood (PLL), to evaluate the extent to which each parameter can be estimated from the available data^9^. Such an approach will inform a maximally informed design of experiments (MIDOE).

Profile Log-Likelihood is a technique to systematically characterize the identifiability of parameters^9,10^. A parameter is generally classified into three categories: identifiable, practically unidentifiable and structurally unidentifiable. Parameters which can be accurately estimated over the entire parameter regime from the available data, are defined as identifiable. Those which can be accurately estimated over a sub-regime are practically unidentifiable and others which cannot be estimated at all are structurally unidentifiable. For an identifiable parameter, the negative Log-Likelihood (numerically equivalent to χ^2^) can be uniquely estimated and conforms to a quadratic profile such that the value of χ^2^ crosses the confidence-threshold (CT) on both sides of the minimum value. However, an unidentifiable parameter deviates from the quadratic profile and cannot be uniquely estimated^3,9,10^.

To determine the extent to which each parameter in the Smad model topology can estimated by fitting to the RFP-Smad2 dynamic data, we evaluated the PLL for each parameter over the entire feasible range (see Methods). First, we created an array of uniformly distributed grid points in log-space over the feasible parameter range. Then we evaluated 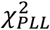 by systematically varying each parameter over the grid points and re-optimizing the others. We implemented the evolutionary strategy, *ISRES+*^20^, for these evaluations at least four times, illustrated as scatter points (**Figure 4**). The average profile for each parameter is visualized as a moving average through the average value of the scatter points (**Figure 4**; black curves). We computed the CT from the distribution of 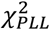 for the Smad_v0 model (**Figure 2A**; colorbar) to be 1.6.

**Figure 4:**
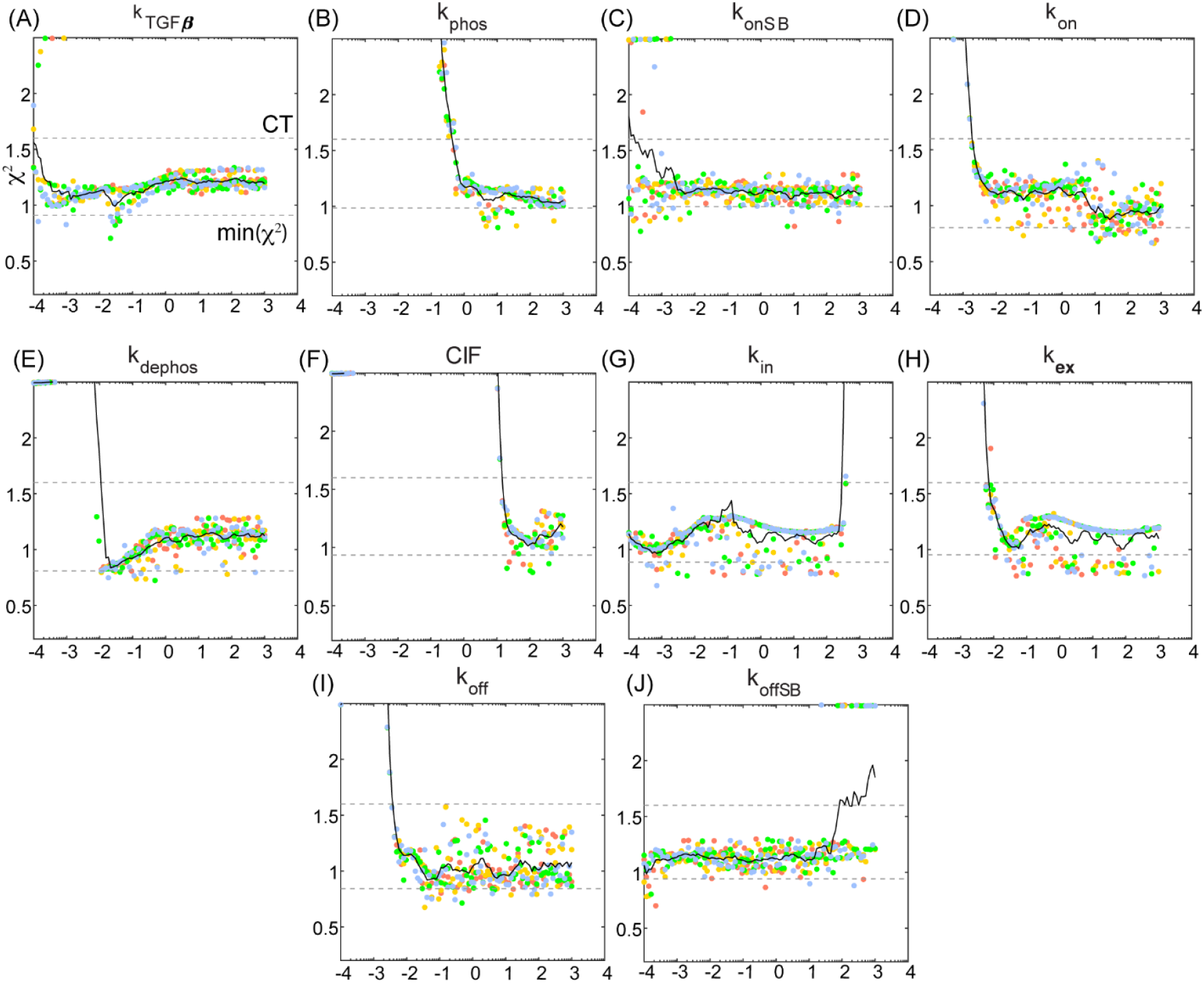
Profile Log-Likelihood analysis on the Smad model calibrated with Smad2 dynamics. **(A-J)** The profile log-likelihood of all the parameters in the Smad model. The (point-wise) confidence threshold was computed as the log-likelihood of the best fit parameter set plus the 99%-quantile of the χ^2^-distribution with degrees of freedom equal to number of parameters. The CT_α_ was calculated to be 1.6.

The ten variable parameters in the Smad model topology can be further classified into two sub-categories. The first category consists of the two *in vitro* parameters, k_onSB_ and k_offSB_, formulated to model the deactivation of the Smad pathway through the addition of an inhibitor (SB-431542). Since the cells were treated with a high-dosage of the inhibitor to deactivate the receptors and shut down the signal transduction, we expect the parameters k_onSB_ and k_offSB_ to be structurally unidentifiable, which is evident through nearly-flat PLL profiles of each (**Figure 4C,J**).

The remaining eight biophysical parameters, which fall into the second category and characterize Smad dynamics *in-vivo*, include rate constants that govern import (k_in_)-export (k_ex_), binding (k_on_)-unbinding (k_off_), and phosphorylation (k_phos_)-dephosphorylation (k_dephos_) rates. Even though the PLL of some of these parameters, including k_on_, k_dephos_, CIF, k_in_, k_ex_ and k_off_, have either one or multiple minima, no parameter in the Smad_v0 is identifiable in the entire range over which it is defined (**Figure 4**).

In addition to determining the identifiability of the parameters in the Smad_v0 model, we can draw several other key inferences to aid experimental design. For instance, we can evaluate the utility of potential experiments such as Fluorescence recovery after Photobleaching (FRAP), which are performed to estimate protein nuclear import (k_in_) and export (k_ex_) rates. The PLL of k_in_ identifies multiple local solutions of which only 10^−3^ min^-1^ is biologically-relevant (i.e., less than 10 min^-1^) (**Figure 4G**). Similarly, of the multiple solutions for k_ex_, only 10^−1^ min^-1^ is biologically-relevant (**Figure 4H**). Since the inferred estimates for k_in_ and k_ex_ from the Smad_v0 model are roughly accurate, performing these experiments would not constrain the parameters and model predictions sufficiently (**Figure 4I-L**).

### Evaluating Candidate Experiments With Profile Log-Likelihood

In the previous section, we rejected FRAP as a candidate experiment through an approximate order of magnitude analysis. In this section, we will utilize the PLL of each parameter to rationally evaluate three other candidate experiments and determine their effectiveness in maximally-constraining model predictions. The three candidate experiments are: (**v1**) measuring the bulk nuclear dynamics of Smad4_total_ upon TGF-β stimulation (**Figure 5A**), (**v2**) tracing the co-localization of Smad2 and Smad4 to obtain bulk nuclear dynamics of all Smad hetero-dimers (**Figure 5B**), and (**v3**) determining particle number and brightness to estimate the stoichiometry of molecular complexes to deconvolve the dynamics of Smad2 dimers and trimers, essentially estimating the dynamics of SC, the Smad trimer, through direct analysis of the experimental data (**Figure 5C**)^24,25^.

**Figure 5:**
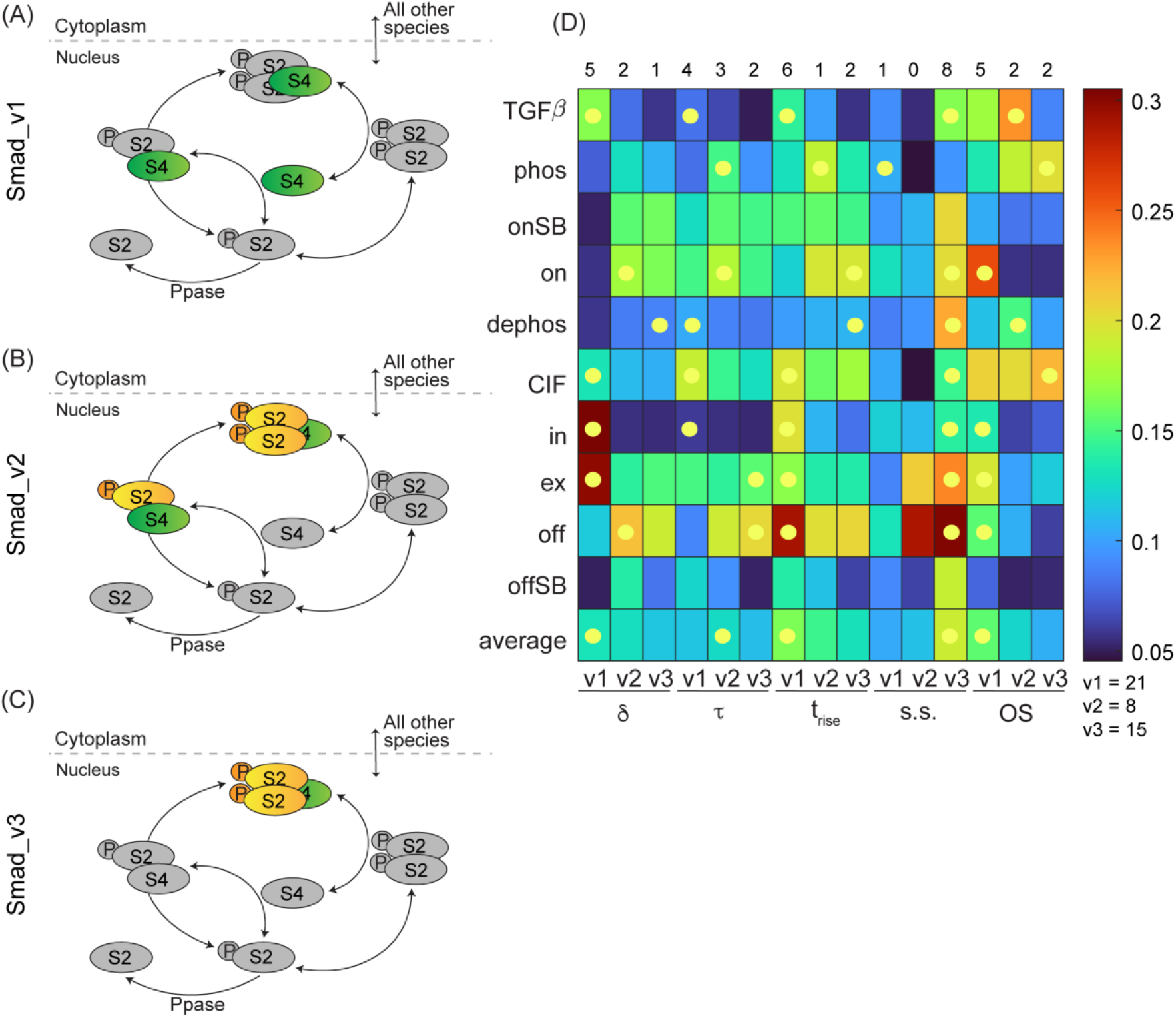
Evaluation of the dynamic characteristics for candidate experimental designs. **(A-C)** Illustration depicting the measurable variable (colored) for the three candidate experiments: total nuclear Smad4 dynamics **(A)**, co-localization of Smad2-Smad4 **(B)** and, estimating the dynamics of the SC **(C). (D)** The standard deviation in dynamic characteristics for the model outcomes with Smad_v0 PLL parameter sets. The yellow marker indicates the experiment with highest variation, which is tallied on top of the heatmap. The sum of the tally for each candidate experiment is shown below the colorbar.

The goal of this approach is to simulate the candidate experiments with parameter combinations obtained in evaluating the PLL for each parameter (below the confidence threshold) and identify those experiments which have the highest variation in their dynamics. The rationale is that, if varying a select parameter results in highly varying predictions for the candidate experiment, then the experimental measurements of that candidate experiment would maximally-constrain the selected parameter. To quantify the variability in the simulated predictions of the candidate experiments, we evaluated their dynamic characteristics using five metrics: dead time (δ), time constant (τ), rise time (t_rise_), steady state (s.s.) and fractional overshoot (OS). We visualized the standard deviation of the estimated metric for each candidate experiment as a heatmap (**Figure 5D**). This approach allows us to evaluate the effectiveness of each experiment in constraining individual parameters as well as the maximally-informative experiment on average. We tallied the effectiveness of each experiment (**Figure 5D**; top row) and identified that the simulated dynamics of experiment (v1) have highest variation and would be maximally-informative in constraining the model predictions.

Note that we proposed these candidate experiments as they are reasonable and typically performed by several Bioengineering Labs. The methods we discussed here could also be applied to evaluate the effectiveness of an alternate set of proposed experiments in calibrating the model, as long as the experimental results could be properly simulated.

### Validating THE Effectiveness OF Proposed Candidate Experiments

The PLL analysis suggests that the candidate experiment (v1) would be maximally-informative in constraining the model parameters and hence predictions, as opposed to experiments (v2) and (v3). Even so, we re-calibrated the Smad_v0 model with the simulated measurements of each of the three experiments (v1, v2 and v3), which resulted in the upgraded Smad_v1, Smad_v2 and Smad_v3 respectively. Although the exact quantitative data that characterize the proposed experiments in v1, v2, and v3 are not currently available in literature, we nevertheless generated approximate dynamics using a combination of simulations and *in vitro* and *in vivo* data^15,19^ from the literature (see Methods). For our purposes, we take these approximated data as ground truth for v1, v2, and v3 (**Figure 6G**, **7C** and **7G**; green).

**Figure 6:**
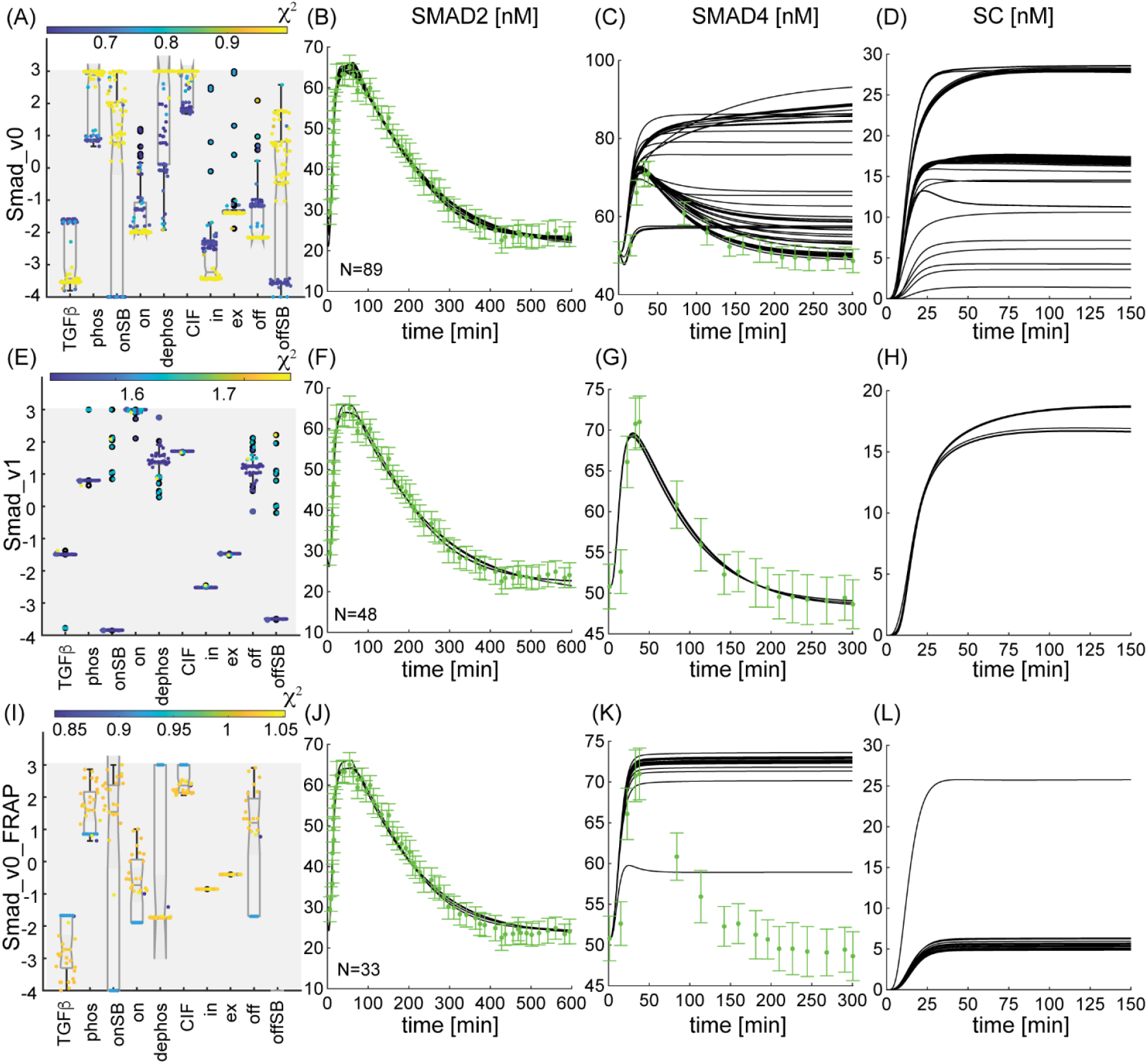
Model fits and predictions for the Smad signaling model calibrated with the Smad2 and Smad4 dynamic data, and the FRAP data. **(A-L)** The inferred parameters **(A,E,I)**, Smad2 dynamics, **(B,F,J)** Smad4 dynamics, **(C,G,K)**, and the predictions of the SC **(D,H,L)** for the Smad model calibrated with RFP-Smad2 data, referred to as “Smad_v0” **(A-D)**, RFP-Smad2 and GFP-Smad4 data, referred to as “Smad_v1” **(E-H)**, or RFP-SMAD2 dynamics and FRAP data **(I-L)**. Black curves represent model fits (B,F,G,J) or model predictions (C,K,D,H,L). Green points represent data (error bars are ±S.D.)^15,19^.

As projected by the PLL analysis, the parameter inferences and predictions are maximally-constrained for Smad_v1 compared to Smad_v2 and Smad_v3 (**Figures 6**-**7**). The Smad_v2 is completely unsuccessful in constraining the predictions of SC and the success of Smad_v3 could be attributed to the fact that we are measuring exactly what we sought to predict. The PLL analysis allowed us to formally evaluate candidate experiments. However, in retrospect, it is evident that the predicted dynamic profiles of Smad4 in Smad_v0 have two distinct behaviors, which may explain why the Smad4 measurements (v1 experiments) are suggested to significantly constrain model predictions.

In addition to the candidate experiments, we also re-calibrated the Smad_v0 with FRAP data (albeit in a different cell-line) and called it Smad_v0_FRAP^26^. As suggested by the PLL analysis, FRAP measurements do not significantly constrain the parameters and predictions (see **Figure 6I-L**).

Finally, we note that the predicted SC dynamics in Smad_v1 (**Figure 6H**) and our “ground truth” dynamics of SC for Smad_v3 (**Figure 7G**) have similar dynamics but distinct steady state levels. While both sets of SC dynamics are predictions, and not direct measurements, we expect the predicted SC dynamics from Smad_v1 to be more accurate, as the Smad_v1 model was calibrated with Smad2 and Smad4 data from the same cell line and set of experiments^15,19^. On the other hand, the “ground truth” SC data from Smad_v3 was generated by using *in vitro* data and simulations. As such, we expect that the discrepancy in the SC steady state between Smad_v1 and Smad_v3 stems from the higher quality data used in Smad_v1. As such, if direct measurements of the SC were performed in the same cell line and under similar conditions as those used to acquire the Smad2 and Smad4 data^19^, we would expect those measurements of SC to align with the predicted dynamics in Smad_v1. Additionally, one set of predictions for SC in Smad_v2 does align with the SC dynamics in Smad_v1, even when the ground truth for the candidate experiment (v2) was approximated from the same *in vitro* data and simulations as (v3).

**Figure 7:**
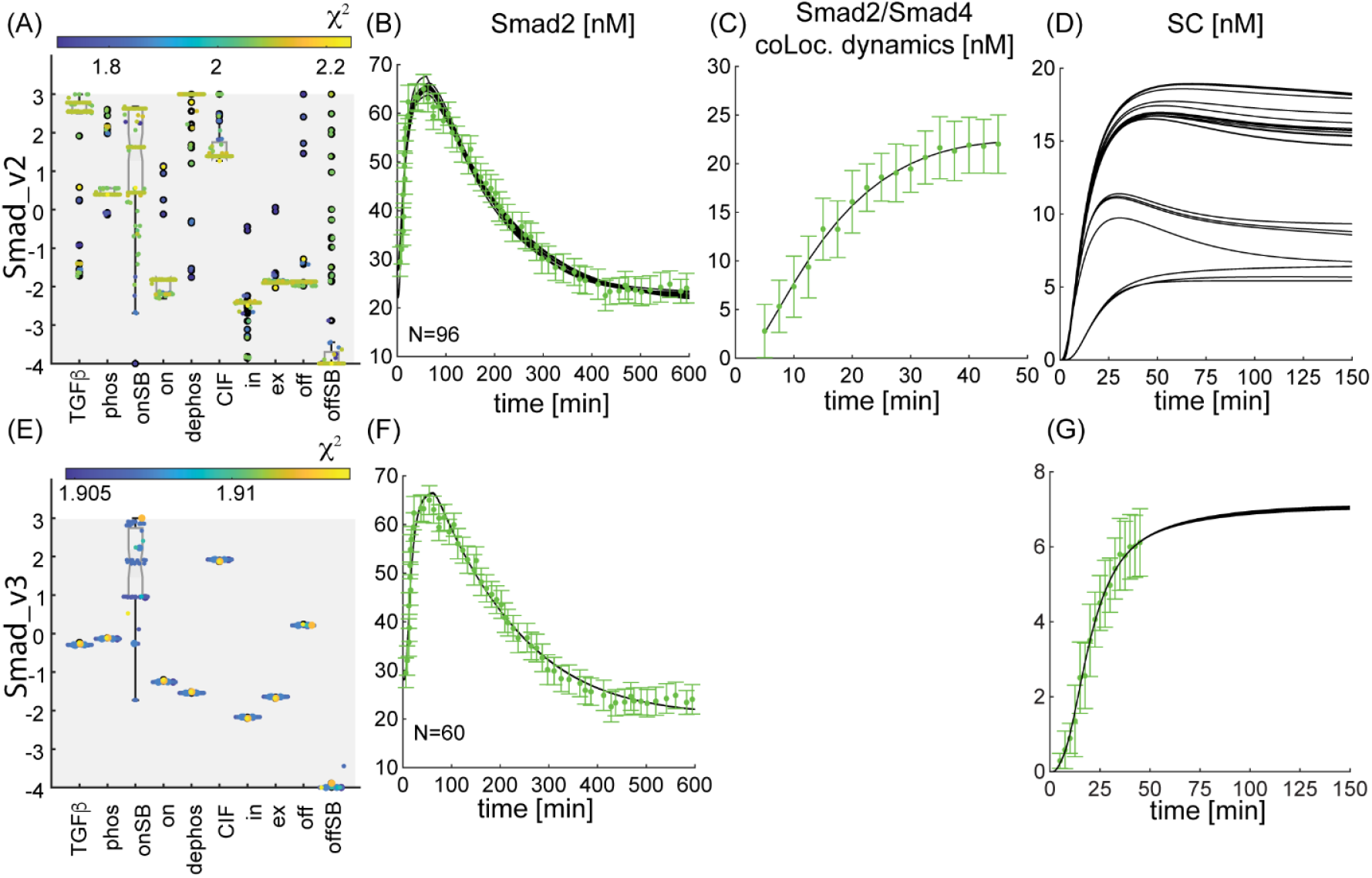
Model fits and predictions for the Smad signaling model calibrated with the Smad2 and Smad2/Smad4 colocalization data, and the SC dynamic data. The inferred parameters **(A,E)**, Smad2 dynamics **(B,F)**, and the predictions of the SC **(D,G)** for the Smad model calibrated with RFP-Smad2 data and Smad2/Smad4 colocalization dynamics **(C)**, referred to as “Smad_v2” **(A-D)** or RFP-Smad2 and SC dynamics **(G)**, referred to as “Smad_v3” **(E-G)**.

### A Less Sloppy Model OF THE Smad Signaling Pathway

As described in the earlier sections, the distribution of inferred parameters for the Smad_v0 spanned across several orders of magnitude, resulting in under-constrained model predictions. By recalibrating the model with maximally-informed experiment, we could constrain the distribution of inferred parameters and the predictions of SC, as indicated by the dynamic characteristics including dead time, time constant, rise time, steady state concentration of SC and fractional overshoot (**Figure 8E**-I). The geometric mean of the eigenvalues of the FIM (gm(λ_FIM_)) of Smad_v1 shifts to the right, indicating that the model is significantly less sloppy and parameter uncertainty is significantly lower (**Figure 8J-K**).

**Figure 8:**
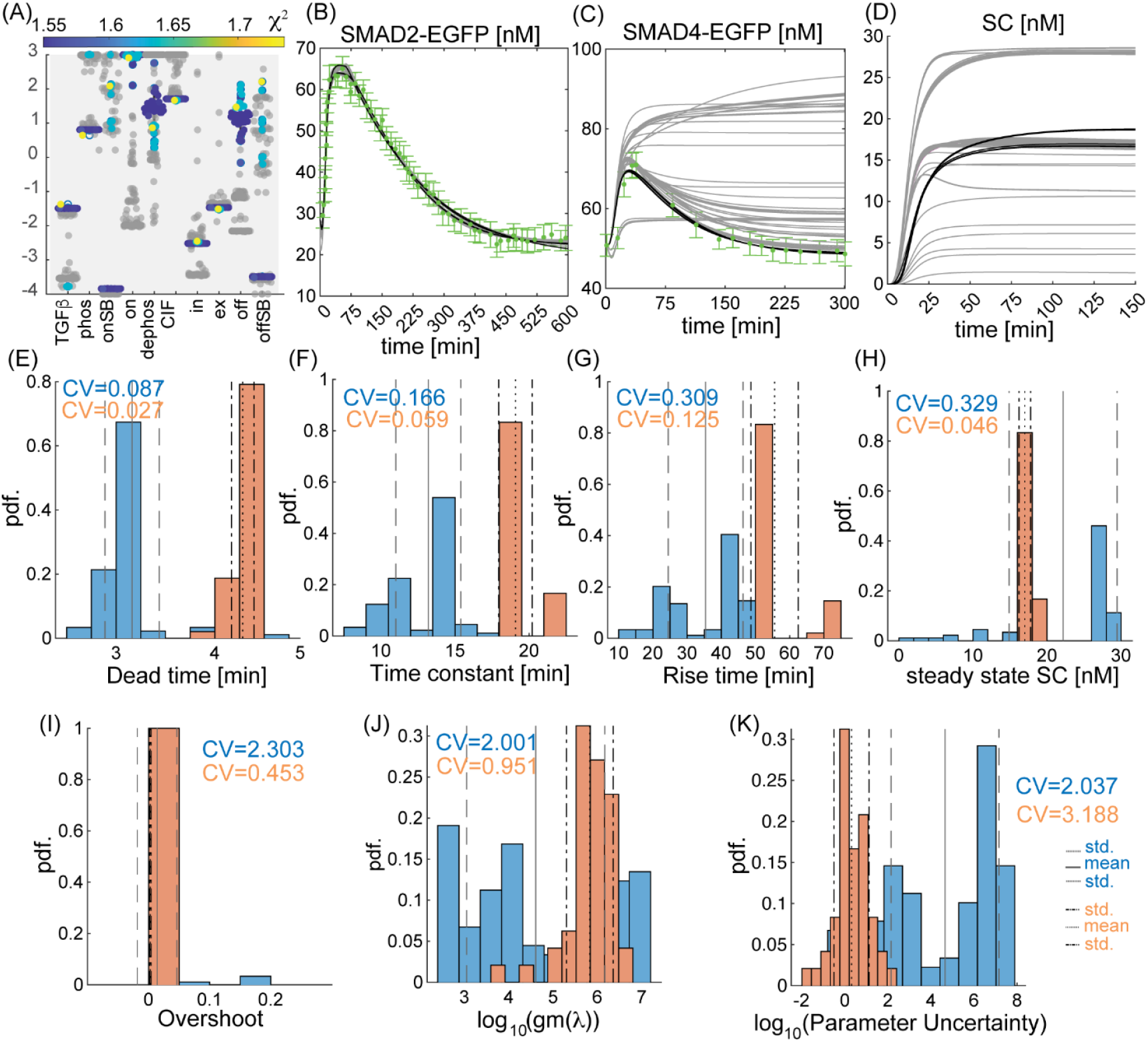
Comparison of the model fits and predictions for the Smad_v0 and Smad_v1. **(A-D)** On comparing the two models (Smad_v0 in gray curves, also in Figure 6A-D; Smad_v1 in black curves, also in Figure 6E-H), the distribution of inferred parameter sets **(A)** for the Smad signaling model calibrated with Smad2 **(B)** and Smad4 **(C)** dynamics relative to the model calibrated on Smad2 dynamics alone results in constrained SC dynamics **(D)**. In (B-D), gray curves represent Smad_v0 model, also in Figure 6A-D, and black curves represent Smad_v1 model, also in Figure 6E-H. Green points represent data (errorbars are ±S.D.)^15,19^**(E-I)** The histograms of the dynamic characteristics of the SC including, dead time **(E)**, time constant **(F)**, rise time **(G)**, steady state concentration of SC **(H)** and fractional overshoot **(I)**; have tight distributions for the Smad_v2 relative to Smad_v1. **(J-K)** Model calibration metrics including geometric mean of the eigenvalues of the FIM **(J)** and parameter uncertainty **(K)** are constrained for the Smad model calibrated on Smad2 and Smad4 information, relative to the Smad model calibrated on Smad2 information only.

We repeated the PLL analysis (as described previously) for Smad_v1 and noted that several parameters, which were unidentifiable for Smad_v0, were now identifiable: k_phos_, k_on_, k_ex_ and k_off_ (**Figure 9**). However, several other parameters, including k_TGFβ_, k_dephos_, CIF and k_in_, remain unidentifiable. One could propose several other experiments to effectively calibrate these and evaluate their effectiveness using the Smad_v1 PLL analysis, but since we have met our goal of constraining the predictions of SC within experimental error, we did not investigate this further.

**Figure 9:**
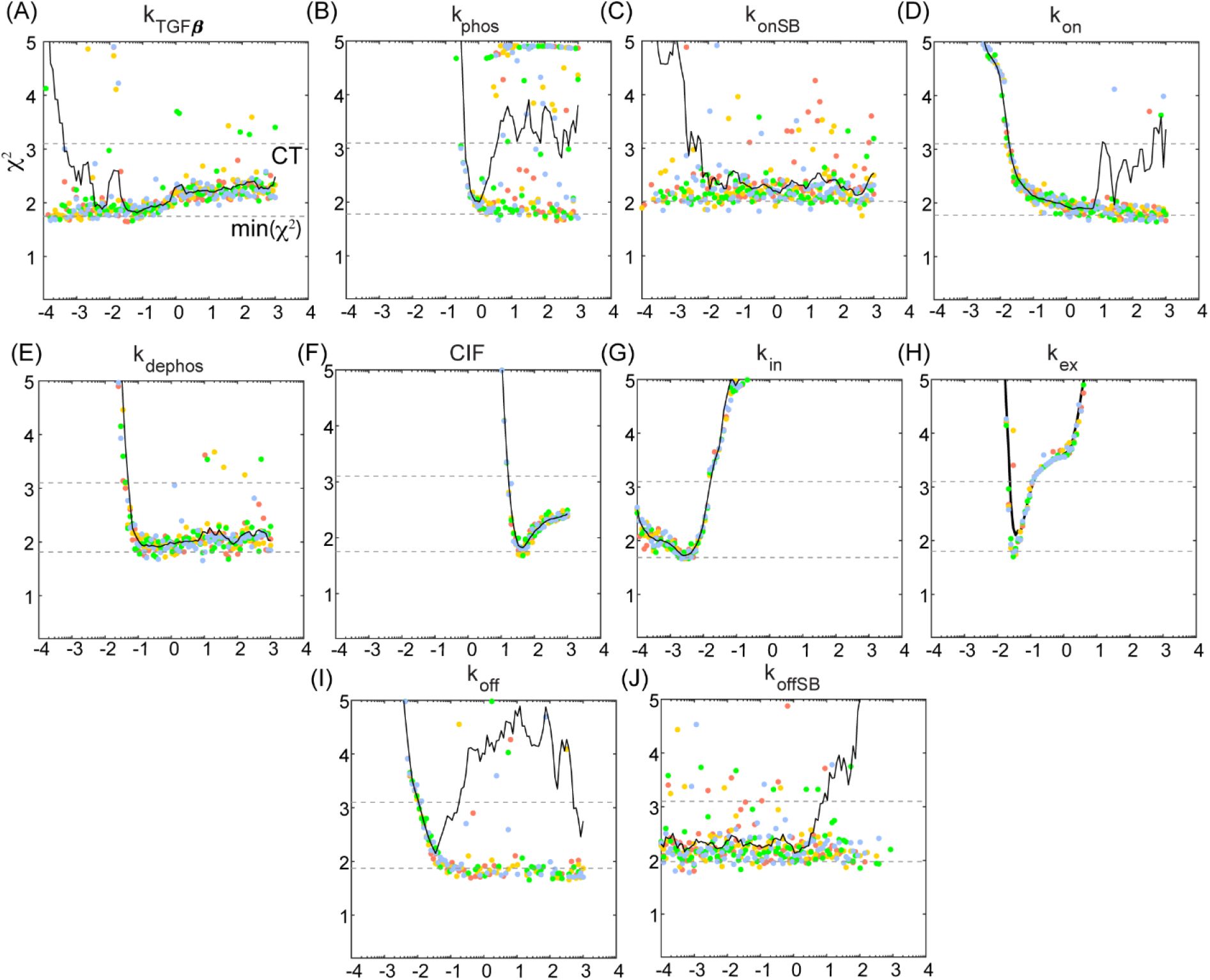
Profile Log-Likelihood analysis on the Smad model calibrated with Smad2 dynamics. **(A-H)** The profile log-likelihood for all the parameters in the Smad model. The (point-wise) confidence threshold was computed as log-likelihood of the best fit parameter set plus the 99%-quantile of the χ^2^-distribution with degrees of freedom equal to number of parameters. The CT_α_ was calculated to be 3.1.

## 4 Discussion

Signal transduction pathways, particularly the TGF-β/Smad signaling pathway, transduce extracellular cues by triggering several complex intracellular interactions to ultimately form a transcription factor and precisely control cellular responses. Due to the emergent dynamics of intracellular Smads, it is challenging to deconvolve measurements and estimate the dynamics of the trimeric signaling complex (SC). One could use an accurate mathematical model to deconvolve the measurements of RFP-tagged Smad2 to account for Smad homo-/hetero dimers and trimers and thereby estimate the dynamics of the transcription factor SC. This approach would require estimating the model parameters by fitting the model to time course data of RFP-Smad2.

However, accurate estimation of model parameters is challenging. Typically, for systems biology models, the experimental data are sparse relative to the number state variables and free model parameters, which impedes accurate parameter estimation. The uncertainties in parameters translate to under-constrained model predictions, such that a large number of “well-fit” parameter sets result in divergent model predictions. Geometrically, this implies that the model manifold, an object of all feasible predictions in high-dimensional data space, is sloppy^27^, which is a common characteristic of systems biology models^7^. In such cases, the eigenvalues of the Fisher Information Matrix (FIM)^2,28^, which is a measure of the amount of information an observable random variable carries about an unknown parameter, spans over several decades. The widths of the axes of the model manifold are inversely proportional to the eigenvalues of FIM, rendering a hierarchy of widths to the shape of the object. Due to the large aspect ratios, the widths range from stiff (narrow) directions to sloppy (wide) directions.

We formally evaluated the sloppiness of an existing predictive model of the TGF-β/Smad signal transduction pathway by generating several “well-fit” parameter sets. These parameters have large uncertainties associated with them (O(θ) ≈ 10^8^) and the eigenvalues of the FIM span over 15 decades. The resulting predictions are under constrained and have highly-varying dynamic characteristics. By reconstructing a pseudo-model manifold around each inferred parameter set, we showed that the model sloppiness is a significant contributor to the variation in the predictions. Further, we computed the Profile Log-Likelihood^9^ (PLL) to assess the extent to which each parameter could be inferred given the model formulation and data — parameter identifiability.

To increase the predictive capabilities of the TGF-β/Smad model, we then used the PLL analysis to propose a maximally-informed design of experiments (MIDOE). This allowed us to project the ability of three candidate experiments in constraining model predictions. We used data from literature to show that PLL-based MIDOE reduces parameter uncertainty and results in constrained model predictions. We also show that PLL analysis helps identify parameters/experiments, such as FRAP, which would not help reduce the variability in model predictions.

With this approach, we show that models in systems biology are sloppy, and several “well-fit” parameters result in highly under constrained model predictions. To accurately predict system behavior, we need to evaluate model calibration and parameter identifiability followed by maximally-informed design of experiments. Such careful analysis would not only improve the predictive power of models in systems biology but also reduce the time and expense of performing non-optimized experiments.

## 5 Methods

### The Smad Signal Transduction Model

#### Model Construction

The model of the Smad pathway was adapted from the Schmierer et al. (2008) model of the pathway fit to EGFP-Smad2 data in HaCat cells^14,15^. In these models, a TGF-β ligand from the family of Transforming Growth Factors - β (TGF-β), binds to its cognate transmembrane receptors resulting in the phosphorylation of the receptor-regulated Smad2 forming PSmad2. PSmad2 interacts with a common mediator Smad, Smad4 to form several hetero- and homomeric Smad complexes. In particular, the trimeric complex (PSmad2)_2_/Smad4 is a transcription factor that regulates gene expression.^11^

However, these models were formulated such that they consisted of EGFP-tagged- and non-tagged Smad2 species. To simplify the model and enable comparison of several candidate experimental designs, our Smad model does not distinguish between EGFP-Smad2 and Smad2.

In our model of the Smad pathway (Supplementary Table **1-2**), monomeric Smad2 and Smad4 are present in the two compartments, the nucleus and the cytoplasm. The activation of the pathway results in the formation of several Smad species including, PSmad2, PSmad2/Smad4, PSmad2/PSmad2 and (PSmad2)_2_/Smad4. These model species can exchange between the nucleus and the cytoplasm. The pathway is deactivated upon the addition of a receptor inhibitor SB-431542. A multiplicative complex import factor (CIF) to the import rate of (PSmad2)_2_/Smad4 denotes enhanced nuclear localization of the complex relative to other Smad complexes.

#### Model Calibration

We utilized the Improved Stochastic Ranking Evolutionary Strategy Plus (ISRES+)^20^ to estimate the ten variable parameters in our model (Supplementary Table **4**). This was achieved by minimizing the sum of squared errors (SSE) between the experimental data and model predictions, normalized by standard deviation.

ISRES+ is a (μ-λ)-based evolutionary algorithm designed to solve global optimization problems using stochastic ranking. It incorporates gradient-based search enhancements, specifically lin-step and newton-step, to refine its search strategy. The algorithm ran for 3000 generations with 400 individuals, using a recombination rate of 0.85. In each generation, lin-step contributed two individuals, while Newton-step contributed one. A full description of the hyperparameters and parameter bounds used in ISRES+ is provided in Supplementary Tables **5** (Config. 1).

The Smad model was calibrated to nuclear RFP-Smad2 dynamics in C2Cl2 cells in response to TGF-β signaling, and later when an external inhibitor of the Smad pathway was introduced (see Supplementary Table **1** for initial conditions). We chose to calibrate the model to RFP-Smad2 data in C2Cl2, since Warmflash et al. (2012) reported live traces of RFP-Smad2 and GFP-Smad4, which we could later use to validate candidate experimental designs. However, Schmierer et al., (2008) data were calibrated onto a concentration scale while Warmflash et al. (2012) reported their data in arbitrary units. Using the EGFP-Smad2 traces (concentration scale) from Schmierer et al. (2008), we approximately calibrated the RFP-Smad2 traces to a nM concentration scale and subsequently the GFP-Smad4 data. The experimental error in the RFP-Smad2 and GFP-Smad4 dynamics were computed as a standard deviation after adding random noise in the range (−5,5) nM to the traces.

The ground truth for the colocalization dynamics of Smad2/Smad4 were approximated from simulated dynamics and relative intensity of phosphorylated Smad2 visualized through a Western Blot. For this, we visualized the dynamics of total phosphorylated Smad for all the parameter sets in Smad_v0. We manually selected the parameter set which most closely reproduced the total phosphorylated Smad from Schmierer et al. (2008) and use it to generate colocalization and (pSmad2)_2_/Smad4 traces to which we added random noise to mimic experimental error.

For the Smad_v1, Smad_v2 and Smad_v3, the objective function was formulated as the sum of two terms: first, the SSE between the Smad2_total_ fits and RFP-Smad2 data; second, the SSE between the corresponding model predictions and the ground truth for each candidate experiment.

### Dynamic Characteristics

The dynamic characteristics for the response are defined below. Note that *y* the response variable, which could be the dynamics of the Smad trimer (SC) or candidate experimental designs. The response variables for the candidate experimental design (v1), (v2) and (v3) are the dynamics of Smad4_total_, colocalization dynamics of Smad2 and Smad4, and dynamics of SC, respectively.

1. Dead time (δ) and Time constant (τ): These metrics were computed by fitting the response (y) to the following function *f* using a MATLAB function lsqcurvefit. The initial guesses for τ were estimated by interpolating onto the time point during which the response was at 37% change from its initial value (y_init_); and the initial guess for δ was 2 min.

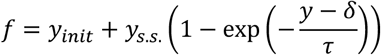
2. Rise time (t_rise_): The rise time is defined as the duration of a response to reach 95% of its steady state value^29^. We calculated the rise time using a MATLAB function stepinfo from the control systems toolbox.
3. Steady State (s.s): Since we ensured that all simulations were run to steady state (dy/dt = 0), the steady state is the last value of the corresponding response.
4. Fractional Overshoot: Relative to the normalized response y_norm_(t) = (y(t) – y_init_)/(y_final_ – y_init_), the overshoot is the larger of zero and max(y_norm_(t) – 1). We calculated the overshoot using a MATLAB function stepinfo from the control systems toolbox.

### Model Calibration Metrics

#### Fisher Information Matrix

The Fisher-Information Matrix (FIM) is a measure of the amount of information an observable random variable carries about an unknown parameter^2,4-6^. It is formulated as a product of the transpose of the Jacobian times Jacobian.

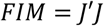

We computed the Jacobian of the Smad model using finite differences, such that each element of the matrix was calculated as below:

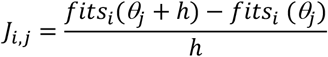

Here *i* ∈ *N*_*fits*_, *j* ∈ *N*_*parameters*_, *N*_*fits*_ denotes the length of the model fits array, and *N*_*parameters*_ is total number of parameters in the model. We set h = 10^−5^,

The variable “fits” refers to the model fits to either the RFP-Smad2 data for Smad_v0; or to the RFP-Smad2 and GFP-Smad4 data for Smad_v1.

#### Parameter Uncertainty

Near the best-fit solution, there is a range of parameter values that, while not optimal, still align with the data within the limits of experimental noise. This region defines the confidence interval for the parameter estimate. The corresponding variation in parameters is defined as parameter uncertainty (PU). This can be approximated as from the FIM^2^.

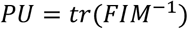

### Psuedo-Model Manifold

The width (w) of the model manifold is inversely proportional to eigenvalues (λ) of the FIM, such that the smallest eigenvalues render the sloppiest widths. With parameter set (*θ*^∗∗^) as its true minimum and a *χ*^2^ value of *χ*^2*\*\**^the i^th^ width of manifold along i^th^ eigenvalue is:

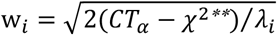

The (point-wise) confidence threshold (CT) is defined as the log-likelihood of the best fit parameter set plus the 99%-quantile of the *χ*^2^-distribution with degrees of freedom equal to number of parameters^3,10^.

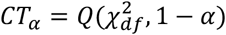

We selected the three sloppiest eigenvalues (*λ*_1_, *λ*_2_, *λ*_3_), and their corresponding eigendirections (*d*_1_, *d*_2_, *d*_3_) to create an ellipsoidal sampling space bound by the confidence threshold (*CT*_*α*_). With the reference parameter set (*θ*^∗^), which has a χ^2^ value of *χ*^2∗^, at the center of the ellipsoid and a confidence threshold of *CT*_*α*_, the k^th^ sampled parameter of the total N_s_ sampled parameters is given by:

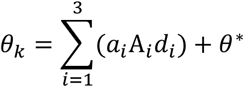

Such that 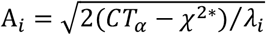 and *a*_*i*_ = −1 + (1 + 1)*rand*(*N*_*s*_, 3). In our sampling, we used N_s_= 1000.

Of all the sampled parameters, negative parameter combinations and those which result in a *χ*^2^ > *CT*_*α*_ were discarded. This is because, for several parameter combinations, the assumption that the pseudo-model manifold geometrically aligns with the actual model manifold does not hold true (**Figure 3D**). All the remaining parameter sets are used to simulate the model predictions and estimate dynamic response characteristics. The coefficient of variation across all the sampled sets results in a single data point in **Figure 3E-F**.

We created the pseudo-model manifold for all the optimal parameter sets (positive-definite) identified for Smad_v0.

### Profile Log-LIKELIHOOD

#### Algorithm

The likelihood represents the joint probability that the independently distributed measurements (*y*) of compound *i* at time point *t*_*j*_, such that each point y_i_ follows a distribution p^3,9^. Mathematically, it is formulated as

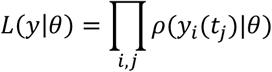

The negative log-likelihood for normally distributed measurement noise corresponds to χ^2^— a measure of agreement between the model predictions and experimental data, and is represented as follows^3,10^:

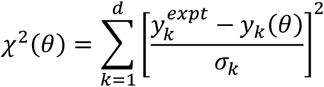

Where *d* denotes the number of data points

To compute the profile of 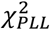 over the entire feasible range i.e. (10^−4^, 10^3^) for each parameter *θ*_*l*=*1*,…,*N*_ we systematically varied the value *θ*_*l*_ over 100 grid points uniformly distributed in logspace. For each value of *θ*_*l*_, we re-optimized all the other free parameters *θ*_*m*≠*l*_ to arrive at a set of parameters for which 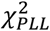 is minimized. We employed an evolutionary algorithm, ISRES+, to determine the optimal set of parameters for each given value of *θ*_*l*_, and repeated this process at least four times for each value of *θ*_*l*_. The hyperparameters for each ISRES+ implementation are listed in Supplementary Table 5 (Config. 2).

To visualize the continuous profile for individual parameters we used robust local regression using weighted linear least squares and a 2^nd^ degree polynomial model of the mean *χ*^2^ over all ISRES+ runs for all 100 values of *θ*_*l*_. This was performed using the “smooth” function in MATLAB with the method set to “moving average” with a window of 5.

We computed the CT for the Smad_v0 and Smad_v1 by calculating the 99% quantile of their *χ*^2^-distribution of all the best fit parameter sets.

## 6 Acknowledgments

Portions of this research were conducted with the advanced computing resources provided by Texas A&M High Performance Research Computing.

This research was supported by NSF grant 2313692 awarded to G.T.R.

## Notes

### Competing Interest Statement

The authors have declared no competing interest.

